# Every Cell Counts: Tomato Root Responses to Nitrogen at Single-Cell Resolution

**DOI:** 10.64898/2026.02.06.704465

**Authors:** Ryan M Patrick, Rajeev Ranjan, Shantha R Sumanasinghe, Philip J SanMiguel, Kranthi Varala, Ying Li

**Author notes:** Correspondence: Ying Li.

## Abstract

The heterogeneity of cell-types in plant roots provides the structural and mechanistic basis for root responses to developmental and environmental signals, including the availability of the major mineral nutrient nitrogen (N) in the soil. To date, single-cell-resolution analyses of transcriptional responses to external nitrogen supply remain limited in plant and crop species. Here, we performed single-cell multiomic profiling of tomato (*Solanum lycopersicum*) roots treated with different nitrogen conditions, generating a high-resolution map of root responses to environmental nitrogen availability. We identified root cell types important for nitrogen response, discovered transcriptional regulators and nitrogen-responsive genes in a cell-type-specific manner, and generated a gene regulatory network across nitrogen conditions that can be used to investigate transcriptional circuits in specific cell types. This single-cell-resolution nitrogen-responsive map of the tomato roots provides a valuable knowledge base for identifying candidate genes to improve nitrogen use efficiency in horticultural crop species.

## INTRODUCTION

Nitrogen (N) is the primary soil mineral nutrient needed by plants (Vidal et al. 2020). Approximately 13 million tons N fertilizer are applied annually in US, accounting for over 60% of chemical fertilizer use (USDA ERS) (Mosheim n.d.). The production of N fertilizer is energy-intensive, and carries long-term environmental impacts (Lehnert, Musselman, and Seefeldt 2021). Ensuring food security and sustainability requires maximizing crop yields while reducing N inputs, which demands a higher nitrogen use efficiency (NUE) in crops.

NUE is a complex agricultural and physiological trait determined by the coordinated actions of multiple biological processes across organs (Q. Liu et al. 2015; Xu, Fan, and Miller 2012). Nitrate (NO_3_^-^), the major form of available N in aerated soil, is absorbed by roots via nitrate transporters (Bian et al. 2025; Léran et al. 2014). Next, NO_3_^-^is either transported to shoots for assimilation, or assimilated directly in the roots, where it is reduced by nitrate reductase and then nitrite reductase (Crawford 1995), and finally assimilated into amino acids via the GS-GOGAT pathway (X. Liu, Hu, and Chu 2022). In addition to being a major nutrient, NO_3_^-^is also an important signaling molecule. The nitrate level is sensed by NPF6.3/NRT1.1 in roots (Léran et al. 2014; Gojon et al. 2011), which initiates a signaling cascade involving calcium signaling and coordinated actions of transcription factor (TFs) (Armijo and Gutiérrez 2017), including the master regulator NLP7 (Marchive et al. 2013). This leads to rapid and widespread reprogramming of transcription of genes encoding transporters, reductases, and aminotransferase enzymes for N assimilation (Varala et al. 2018; R. Wang et al. 2003).

The roots play an essential role in N sensing, transport, and, in some cases, assimilation. The roots display great developmental plasticity in response to N variability, with enhanced lateral root growth in N-rich patches to maximize N uptake (Ruffel et al. 2011; P. Yu, Hochholdinger, and Li 2015). Prior molecular investigations of root responses were primarily performed at the whole organ level (Ruffel et al. 2011; Varala et al. 2018; R. Wang et al. 2003). However, the root is a complex organ consisted of multiple tissues and cell types with distinct functions and molecular signatures (Birnbaum et al. 2003). For example, the primary nitrate transporter NRT1.1 is expressed preferentially in root tips, in agreement with its role of N sensing (Krouk, Lacombe, et al. 2010); meanwhile, genes involved in lateral root initiation in response to local N supply are regulated in the pericycle cells (Ying Zhang et al. 2023). Such genes with cell-type-specific N responses could be overlooked in whole-organ studies. To address this, research on cell-type-specific N responses has been performed in Arabidopsis model (Lhamo and Luan 2021; Gifford et al. 2008), which provided valuable baseline insights. However, Arabidopsis evolved in natural environment where N limitation is common, whereas crop cultivars have been selected and bred in N-supplied soil throughout human agricultural history, therefore, the molecular circuits underlying their N responses could differ significantly. Moreover, research of Arabidopsis roots are often performed in agar plates at seedling stage, while certain cell layers, especially those related to secondary growth, are not observed at this early developmental stage (Wunderling et al. 2018). Furthermore, crop plants may have additional cell layers, such as the exodermis, that are absent in Arabidopsis (Kajala et al. 2021).

Recent studies have begun to fill these gaps by extending cell-specific molecular investigations to crop species, such as wheat (Hai et al. 2025). In addition to cereal models, the horticultural and fruit crop model tomato (*Solanum lycopersicum*), represents a great crop system for nutrient study, because: (*i*) it is the most important fruit vegetable crop, accounting for ∼2 billion dollars in annual farm cash receipt (USDA ERS “*Farm Income and Wealth Statistics*”) and producing ∼100 million tons fresh fruits that serve as rich source of vitamins, minerals, and antioxidants (FAO stats); (*ii*) tomato has a high-quality genome assembly and is highly amendable to genetic modification (Feng et al. 2013), making it ideal for molecular genetics research; and (*iii*) commercial production of tomato relies significantly on greenhouses and hydroponic systems, which are uniquely well-suited for analyzing roots and nutrients. The molecular investigation of N response in tomato was limited (Y. H. Wang, Garvin, and Kochian 2001) but has gained momentum in recent years (Bian et al. 2025; Julian, Patrick, and Li 2023; Bvindi et al. 2022; Sunseri et al. 2023), revealing the genetic basis underlying NUE variation (Sunseri et al. 2023), regulatory role of chromatin modification (Julian, Patrick, and Li 2023; Bvindi et al. 2022), and conservation and difference between Arabidopsis and tomato N responsive circuits (Bian et al. 2025). However, to date, the N response has been investigated mostly at the whole organ level in tomato, while a cell-type-specific study is lacking.

Recently, single cell transcriptomics technology has greatly improved the spatial resolution of cell-type-specific molecular investigation (Hai et al. 2025; Cantó-Pastor et al. 2024; Shahan, Nolan, and Benfey 2021; Shahan et al. 2022). Moreover, single cell approaches could generate massive multi-dimensional functional genomics data, including transcriptomes via RNA-seq and chromatin accessibility profiles via ATAC-seq (Thibivilliers et al. 2023), which provided a comprehensive data set for inferring gene regulatory networks. Complex traits such as NUE are often determined by multiple genes interacting together, collectively represented as gene regulatory networks (GRNs). N-responsive GRNs have been thoroughly studied (Bian et al. 2025; Varala et al. 2018; Brooks et al. 2019; Gutiérrez et al. 2008; Krouk, Mirowski, et al. 2010), fueled by the advance of sequencing technology and computational tools. However, these analyses have usually been performed using aggregate data from whole plant or organ level samples, obscuring the unique contribution of individual cells or cell-types to N response.

In this study, we applied single-cell multiomic profiling to tomato roots under different environmental N supply conditions and generated a high-resolution map of root responses to N. We identified root cell types particularly important for N response, discovered transcriptional regulators and responsive genes in a cell-type-specific manner, and inferred the gene regulatory networks they mediate.

## METHODS

### Plant growth conditions

The nitrogen treatments were performed following a protocol previously published by our lab (Julian, Patrick, and Li 2023). In detail, tomato (*Solanum lycopersicum* cv. M82) seeds were surface sterilized and germinated on ½ MS agar plates in a growth chamber for eight days (16-hr day at 24°C with 100 µmol/s/m^2^ light and 8-hr night at 20°C). Seedlings were then transferred to a hydroponic system with +N growth media (Julian, Patrick, and Li 2023; Y. H. Wang, Garvin, and Kochian 2001) with two hours aeration per day for two weeks. Plants were then transferred to -N media (Julian, Patrick, and Li 2023; Y. H. Wang, Garvin, and Kochian 2001) for 4 days. The seedlings were either treated to resupply with +N media or continued starvation in freshly supplied -N media for 6 hours before sampling.

### Single cell multiomic library construction and sequencing

Nuclei isolation procedures were adapted from previous publications (S. B. Thibivilliers, Anderson, and Libault 2021; S. Thibivilliers, Anderson, and Libault 2020; Nobori et al. 2023). Specifically, for each biological replicate, whole roots from two plants (with a total weight of ∼1.6g) were chopped by razor blade on a petri dish in 3 mL of NIB medium (S. B. Thibivilliers, Anderson, and Libault 2021) for 5 minutes in cold room conditions to form a fine slurry. The slurry was incubated in the cold room for 5 minutes with gentle platform shaking and then successively strained through 40 mm and 30 mm strainers (prewet with 0.5 mL NIB medium) into a 50 mL conical tube on ice. The plate was carefully rinsed with 0.5 mL of NIB medium to collect remaining nuclei. The slurry was centrifuged for 3 min at 50 x g, 4°C to pellet debris. Supernatant was moved to a fresh 50 mL conical tube and centrifuged again for 5 min at 500 x g, 4°C to pellet nuclei. Supernatant was discarded and the nuclei were resuspended in 1 mL NIB medium.

To permeabilize the nuclei, 0.02% NP-40 (Surfact-Amps Detergent Solution, Thermo Fisher) was added to the 1 mL of NIB medium with nuclei suspension to reach a final detergent concentration of 0.01%. Nuclei were incubated with gentle rocking in the cold room for 3 min and then centrifuged for 5 min at 500 x g, 4°C to pellet nuclei. Supernatant was discarded and to remove residual detergent the nuclei were carefully resuspended in 5 mL of PBS+ medium (S. B. Thibivilliers, Anderson, and Libault 2021) using wide bore tips. Nuclei were again centrifuged for 5 min at 500 x g at 4°C to pellet, with supernatant discarded. Nuclei were resuspended in 500 µL of 1x nuclei buffer (10x Genomics, PN-2000207) with 1:40 Protector RNase Inhibitor (Millipore Sigma) and 1 mM DTT. Resuspended nuclei were filtered to remove clumps using a 40 mm Flowmi cell strainer pipette tip (Bel-Art) and then centrifuged for 5 min at 500 x g, 4°C. Supernatant was discarded and nuclei were resuspended in 50 µL of nuclei buffer with RNase inhibitor and DTT.

A sample of prepared nuclei was propidium iodide-stained and imaged with a Nikon Eclipse Ti-S to provide an estimate of nuclei quantity and quality. Images of five large square blocks of the hemocytometer (four corners and one center) were taken, and all nuclei of each large square were counted. Nuclei density was estimated using the formula: (total nuclei counted/5) x10^4^ x dilution factor = nuclei/ml. To attempt to recover 10,000 nuclei per sample, ∼16,100 nuclei were loaded into the Chromium Controller at the Purdue Bindley Genomics Core following the protocol of Chromium Next GEM Single Cell Multiome ATAC + Gene Expression Reagent Kit A (10x Genomics Inc) to generate sequencing libraries. The sequencing of both library types (ssRNA-Seq and ssATAC-Seq) concurrently using four Aviti (Element Biosciences) 2x75 High cassettes was undertaken using the read structure: I1:10, I2:16, R1:50, R2:90, to generate 3.6 billion R2 reads to average 456 million reads per library.

### Data processing

Sequencing reads for ssRNA-Seq and ssATAC-Seq were aligned to the *Solanum lycopersicum* v4.0 genome using Cell Ranger ARC software (10X Genomics) patched for processing AVITI data. SeuratObjects (Hao et al. 2024) were then generated from the filtered feature matrix data produced by the 10X genomics Cell Ranger software. To remove low quality nuclei and possible multiplets, cells were further filtered in Seurat based on assigned feature depth, keeping those with RNA counts >200 and <8001, and ATAC counts >500 but <6000. The SeuratObject data was log normalized and scaled, PCA was run with the top 2000 variable features, and initial clustering of cells was done using 20 dimensions and a resolution of 0.6. The four SeuratObjects (2 replicates each of 2 conditions) were then combined using FindIntegrationAnchors function, utilizing canonical correlation analysis, and using 2000 anchors for integration. The data was then split back into the four individual replicates, retaining the anchor information, and imported into Monocle3 as CellDataSet objects (J. Cao et al. 2019). The four objects were preprocessed with a dimensionality of 50, merged using batch correction alignment, and the 2000 anchor genes were used to cluster the cells with resolution 3e-4. The merged, aligned, and clustered object was then moved back to Seurat for analysis of gene expression, maintaining the projection and cluster assignments from Monocle3.

### Cell type assignment and differential gene testing

Cell-type identities for the 26 clusters were preferentially assigned using previously validated root marker genes from tomato (Cantó-Pastor et al. 2024) and Arabidopsis (Shahan et al. 2022) with additional support from orthologous marker gene groups identified in a large cross-species study (Chau et al. 2025). Relevant specialized literature was utilized as appropriate for support or clarification of cell-type assignment. Differential gene expression testing between -N and +N conditions was performed for individual clusters, for groupings of epidermal, stele, and cortex cells, and for the entire dataset using the FindMarkers function in Seurat (Hao et al. 2024) with a significance threshold of FDR < 0.05, fold-change ≥ 2. Differentially expressed genes were then assigned to one of three categories: (*i*) ‘Broad’ genes were DEGs across more than one cell-type, excepting related cell-types that were part of a tissue level grouping (e.g. epidermis or stele); (*ii*) ‘Type-specific’ genes were DEGs in only one cell or tissue type and subject to an additional condition: for up-regulated genes, the gene expression level was highest in +N condition for that cell-type or tissue; for down-regulated genes the gene expression was either the lowest in +N condition for that cell-type or tissue or was highest in the -N control indicated it was highly expressed in that cell-type/tissue before nitrogen resupply; (*iii*) ‘Type-modulated’ genes were DEGs in only one cell or tissue type, but were not type-specific in nature (that is, they generally had stronger or weaker expression levels in other cell or tissue types but in those cell-types did not meet criteria for significant change in expression). Enriched GO terms were determined using an in-house enrichment testing pipeline previously developed for use in tomato (Julian, Patrick, and Li 2023).

### Gene regulatory network inference

A tomato root gene regulatory network (GRN) was generated from the matrix of single cell gene expression data encompassing all observations across -N and +N conditions, using GRNBoost2 (Moerman et al. 2019) with a previously identified list of tomato TF genes as regulators (Julian, Patrick, and Li 2023); the top 10th percentile of predicted edges was retained. To test for enrichment of various sets of genes among predicted TF-to-target relationships in the GRN, an in-house python script was utilized which performed a series of hypergeometric tests together with multiple testing correction (Benjamini and Hochberg 1995), with an FDR cutoff of 0.05 and a further four-fold cutoff for enrichment level.

## RESULTS

### Mapping cell and tissue types in single cell data collected from whole root systems of tomato seedlings

To investigate genome-wide regulatory responses to N availability at the single cell level, we employed the Chromium Next GEM Single Cell Multiome kit (10X Genomics, CA) to measure gene expression (by scRNA-Seq) and chromatin accessibility (by scATAC-Seq) wherein it is possible to generate both profiles from the same nucleus identified by a unique barcode (Fig. 1A). Briefly, three-week-old hydroponically grown tomato plants (*Solanum lycopersicum* cultivar M82), following four days of N-starvation treatment, were treated with: (*i*) +N: a N supply of 2.8 mM NO_3_^-^, or (*ii*) -N: continued N-starvation of 0 mM NO_3_^-^. After six hours of N treatments, nuclei were isolated from whole root systems of a pool of two individual seedlings experiencing either N-deplete (-N) or N resupplied (+N) condition. Next, transcriptome and chromatin accessibility for each nucleus was profiled using the Chromium NextGEM single cell multiome kit (10x Genomics Inc) at the Purdue Genomics Core Facility (see methods for details). The nuclei preparation, sequencing library construction, and sequencing were done for two independent biological replicates of each N condition.

**Figure 1.**
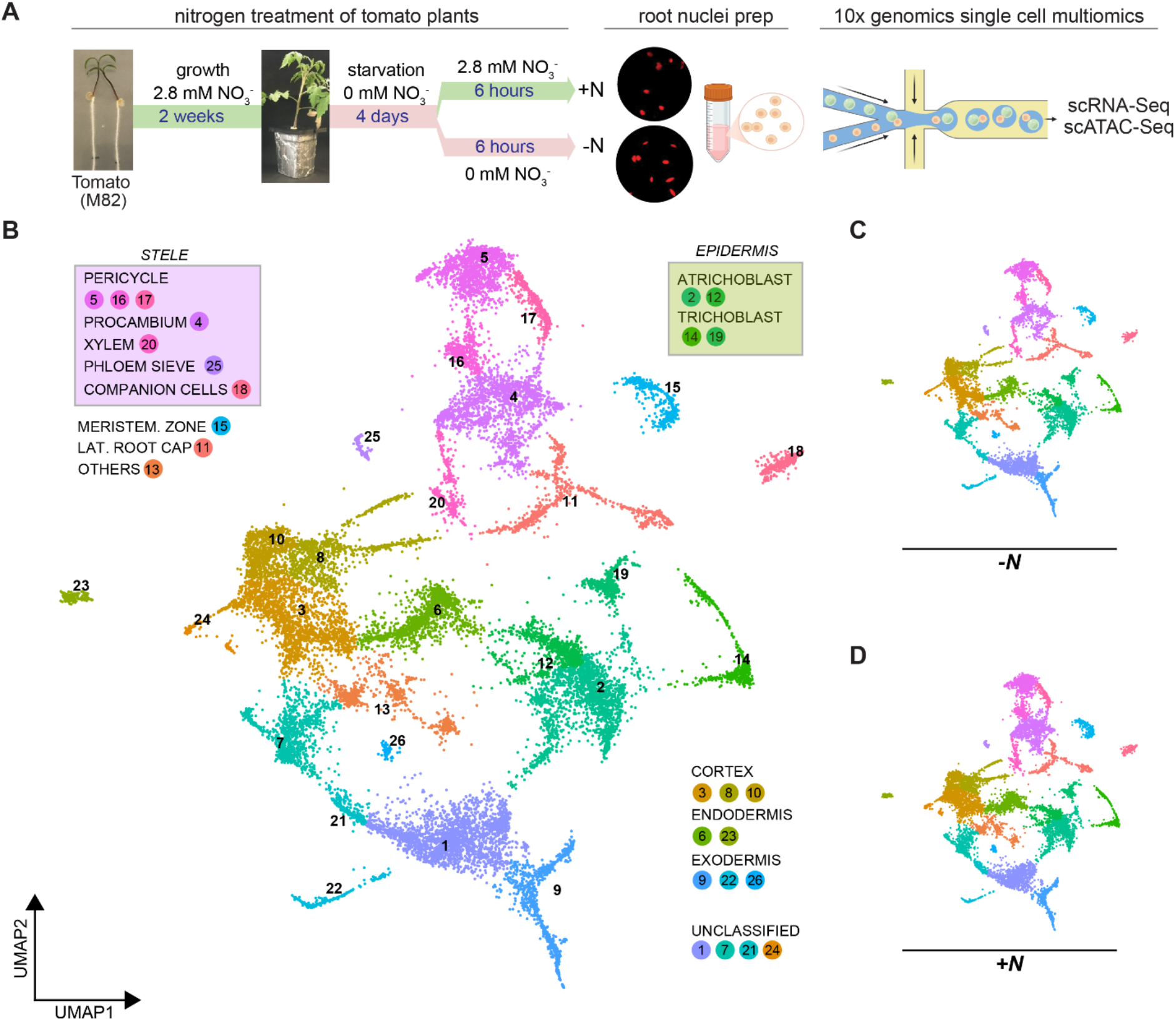
(A) Workflow for tomato single cell experiment. (B) Projection of single cell data across two replicates in two conditions (-N and +N) making up 26 clusters which were assigned cell type identities and additionally larger groupings for stele and epidermal tissues. (C-D) Data from the two N treatment conditions separately shows a similar distribution across cell types.

Following data processing, alignment, and filtering using CellRanger (10x Genomics Inc) and Seurat (Hao et al. 2024), 21,196 root cell nuclei across two biological replicates and both N conditions were clustered according to their gene expression profiles with batch correction in Monocle3 (J. Cao et al. 2019). Among the 21,196 root cells, 26 cell-type clusters were observed, with some being constituents of higer order tissue groupings (Fig. 1B). Cell-type identities for the clusters were assigned using known marker genes, starting with assignments made in a previous tomato root scRNA-Seq investigation (Cantó-Pastor et al. 2024), additionally utilizing root cell atlases developed in Arabidopsis (Shahan et al. 2022) and further bolstered by a broad identification of orthologous marker gene groups from data integrated across 15 plant species (Chau et al. 2025). Using these resources, it was possible to assign cell-type identities to most of the observed clusters.

Epidermal tissue is present as cells forming four clusters (Fig. 1B): clusters 2 and 12 being atrichoblast cells (S. Fig 2) with closely related gene marker patterns (S. Fig. 1), and clusters 14 and 19 being trichoblast cells (S. Fig 2). The trichoblast cells notably diverge in expression of orthologs of certain root hair specific regulators (Shibata and Sugimoto 2019): for example, *SlLRL1a (Solyc12g010170)* and *SlRSL2-like (Solyc12g088380)* are specifically expressed in cluster 14 (S. Fig. 2).

The constituent cell types which make up the stele as a tissue distribute in proximity according to their gene expression profiles in the UMAP (Fig. 1B): the largest group being the procambium at cluster 4, with the pericycle present in one larger cluster, cluster 5, and two smaller clusters, cluster 16 and 17, while xylem is represented in cluster 20 (S. Fig 3). Clusters 18 and 25 represent two distinct distributions of phloem cells marked by *SlAPL* (*Solyc12g017370*.*3*): prior reports of root phloem poll cell development (Otero et al. 2022; Miyashima et al. 2019) were used to identify markers for phloem sieve elements in cluster 25 and the companion cells in cluster 18 (S. Fig 3D).

The cortex was found to have less conserved markers when compared with the previous study in tomato. However, clusters 8 and 10, which have overlapping marker sets with cluster 3 (S. Fig. 1), are enriched in cortex markers as identified from cross-species orthology (S. Fig 4). *SlEXO1 (Solyc09g011120)*, a transcriptional regulator expressed in the inner cortical layer in tomato (Manzano et al. 2025), is present in these three clusters, with the highest expression in cluster 8; the cortex-associated transcription factor *SlJKD (Solyc10g084180)* (Hassan, Scheres, and Blilou 2010; Liang et al. 2024) is also expressed in these clusters (as well as cluster 7), with highest expression found in cluster 3. Together, evidence supports these clusters 3, 8, and 10 as being types of cortical cells.

The endodermis is present across two clusters, one larger, cluster 6, and one relatively small, cluster 23. While endodermal markers (S. Fig 5) are enriched across both, including the cell-type-specific endodermal regulator *SlMYB36* (*Solyc07g006750*), Casparian strip markers which are associated with lignification (Manzano et al. 2025) are highly expressed only in the cluster 23 grouping.

Cluster 9 and the smaller clusters 22 and 26 were identified as exodermal cell types. Markers for developing and suberizing exodermal cells in the tomato root tip were previously characterized in detail (Cantó-Pastor et al. 2024) and are present in cluster 22 and 26 (S. Fig. 6B); however, *SlMYB92* (*Solyc05g051550*), an important regulator associated with developing exodermis, is expressed solely in cluster 22, likely identifying this cluster as active young exodermal cells reported previously in root tips (Cantó-Pastor et al. 2024), while cluster 26 showed strong expression of many genes associated with lignification (*e*.*g. SlPELPK*/*Solyc04g014290* and *SlLAC12-like*/*Solyc05g052370*), indicating that this cluster likely corresponds to a more mature type of exodermal cells (S. Fig 6). Markers for cluster 9 strongly overlap with those identified as exodermis-associated based on orthologous gene markers (Chau et al. 2025) (S. Fig 6A), however, the previously reported exodermis markers (Cantó-Pastor et al. 2024) are not expressed in this cluster. Therefore, one possibility is that this cluster 9 represents an exodermal cell population not present in the previous study (Cantó-Pastor et al. 2024), but present in whole root systems. Two branches are present in cluster 9, which can be identified as having partly overlapping and partly unique sets of expressed genes; for example, *SlMYB68b* (*Solyc06g074910*) is expressed in one branch while *SlWRKY45* (*Solyc08g067360*) is preferentially expressed in the other (S. Fig 6).

Cluster 15 was clearly identifiable as meristematic zone cells (S. Fig 7A) based on previous reported marker genes from tomato (Cantó-Pastor et al. 2024) and Arabidopsis (Komaki and Schnittger 2017; Criqui et al. 2002). Specific markers for the lateral root cap (Bennett et al. 2010) are present in cluster 11 (S. Fig 7B), which may contain other closely related cell types. Cluster 13 is made up of several subclusters with distinct marker populations and may represent stem cells or intermediates in differentiation, and some cells express genes associated with the root cap (Chaiwanon and Wang 2015) and secondary growth (Woerlen et al. 2017; Ye et al. 2021) (S. Fig. 7C).

Finally, four remaining clusters were not possible to clearly assign origins based on available information. Cluster 24 shows a small group of cells with highly divergent gene expression patterns and poor overlap with known root cell marker genes. Clusters 1, 7, and 21 express overlapping sets of markers (S. Fig 1 & 8A), and cluster 1 and 21 gene expression patterns also partly overlap with cluster 9 (putative exodermal cells) (S. Fig 1 & 8A). Notably, *SlMYB68* (*Solyc11g069030*), a member of a group of transcriptional regulators of suberin patterning in Arabidopsis (Manzano et al. 2025; Kraska et al. 2025), is expressed across both cluster 1 and the putative exodermal cluster 9 (S. Fig. 8B), making it an attractive candidate for further investigation of suberization in maturing roots. Another recently described exodermal regulator of suberin deposition in tomato, *SlMYB41* (*Solyc02g079280*), is expressed primarily in cluster 1 (S. Fig. 8C). Given these observations, in the context of this present study which sampled whole root systems including extensive lateral root networks, clusters 1, 7 and 21 may represent a population of mature or lateral root exodermal/cortex cell types which have not been extensively observed in previous studies.

### Identifying root genes involved in either broad or cell-type-specific responses to nitrogen

Across the two biological replicates, 8,748 nuclei were profiled under the -N control condition, and 12,448 nuclei were profiled under the +N condition. Similar clustering by expression patterns was observed between the two N treatments (Fig. 1C-D), indicating consistency in cell identity between the samples. Genes differentially expressed (FDR < 0.05, fold-change ≥ 2) in response to N resupply were then determined by comparisons between the +N condition and the -N control at the whole root level (across all clusters), at the tissue and broader cell-type level groupings (e.g. epidermis, stele, cortex, as shown in Fig. 1B), as well as at individual cluster levels. We observed that differential expression testing in Seurat was very sensitive to cluster size, therefore, to reduce false positives, we also considered percent expression by cluster to increase stringency. Overall, we detected that 1,900 genes are up-regulated in at least one comparison in response to +N treatment, while 962 genes are down-regulated.

We were interested in distinguishing gene expression changes that are broadly shared across many root tissues and cell types from those that are tissue- or cell-type-specific. To do this, DEGs were classified in three ways: broad (whose expression levels significantly changing across multiple cell or tissue types), type-modulated (significantly changing in one cell/tissue type but not specifically expressed in one cell/tissue type), or type-specific (significantly changing and primarily expressed in one tissue or cell-type).

Using this approach, among the 1,900 up-regulated genes, 940 were broadly differentially expressed, 659 showed type-modulated differential expression, and 301 genes were tissue- or cell-type-specific DEGs. Among the 962 down-regulated genes, the numbers were 609 in the broad category, 164 in the type-modulated category, and 189 were type-specific. When broken down by tissue and cell type, the cortex appears to have the highest number of genes that are specifically up-regulated in response to nitrogen (Fig. 2A), and despite being a relatively small portion of cells in the overall root system, the exodermis has the second highest number of genes that are significantly up-regulated and specific to that cell type (Fig. 2A). In the direction of down-regulation, the epidermis has the most type-specifically regulated DEGs followed by the stele and then the cortex (Fig. 2A).

**Figure 2.**
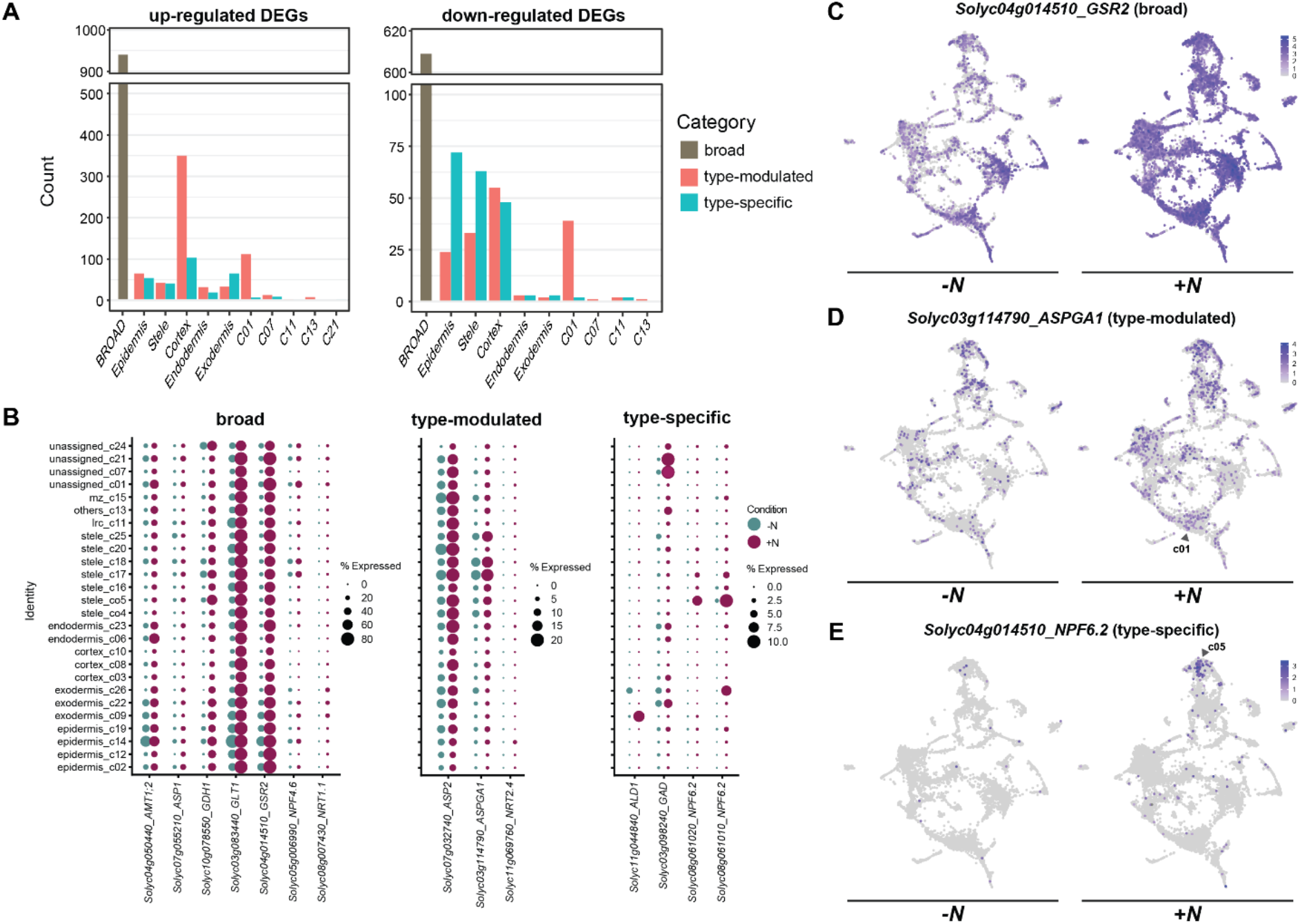
(A) Number of DEGs either broadly up-regulated or down-regulated or showing type-modulated/type-specific regulation by tissue or cell type identity. (B) Dot plots showing expression across the clusters for up-regulated N-responsive DEGs involved in N metabolism and transport across broad, type-modulated and type-specific categories. (C-E) Maps of gene expression for exemplary up-regulated DEGs in each category. Color represents scaled expression level.

These results were compared to our previously published study of bulk RNA-Seq of tomato roots treated with the same +N vs -N conditions for the same length of time (six hours) (Julian, Patrick, and Li 2023).

There was a significant overlap with our previous report – over a quarter of scRNA-Seq DEGs were identified in the previous experiment via bulk RNA-seq (S. Fig 9A-B). Interestingly, there were differences by category: about 36% of broad DEGs identified in the current single cell study were also detected in bulk RNA-seq, but only 18% of the type-modulated and type-specific DEGs were identified by bulk RNA-Seq.

The root DEGs were also overlapped with genes putatively involved in N assimilation and transport based on homology to Arabidopsis genes (Para et al. 2014; Julian, Patrick, and Li 2023). Among these 80 putative N-relevant genes, 14 were up-regulated in the roots, with 7 showing broad responsiveness, 3 showing type-modulated response, and 4 showing type-specific response. Broad up-regulation was noted for several genes involved in N assimilation, as glutamate synthase (*GLT1, Solyc03g083440*) and glutamine synthetase (*GSR2, Solyc04g014510*) were highly induced across most cell types, as well as nitrate transporters (Fig 2B-C). Aspartate aminotransferase (*ASP2, Solyc07g032740*) showed type-modulated up-regulation, in that it is expressed broadly across tissue and cell types, but N-responsive only in the endodermis (Fig. 2B). Among genes with type-specific responses, two copies encoding nitrate transporter (*NPF6*.*2, Solyc08g061010/Solyc08g061020*) were identified to have highly specific induction in the pericycle of the stele (Fig. 2B & E). For N-relevant genes which were down-regulated in response to N supply, 8 showed broad responses while 3 were type-modulated (S. Fig 9C). The broadly down-regulated genes were again involved in nitrogen assimilation and transport, but also included nitrate reductase (*NIA2, Solyc11g013810*) and nitrite reductase (*NIR1, Solyc01g108630*); the three type-modulated genes were all putative N transporters.

Root DEGs were also compared against lists of TFs known to regulate N response, including members of the NIN-like protein family (NLP), HRS1 homolog family (HHO), and lateral boundary domain family (LBD), and TGA transcription factors which have known roles in N response (Julian, Patrick, and Li 2023; Alvarez et al. 2014). Four members of these TF families were up-regulated in the root data, while six were down-regulated (Fig. 3A). Broad up-regulation was observed for *HHO2* (*Solyc05g009720*) (Fig. 3B) while broad down-regulation was observed for *NLP2* (*Solyc01g11219*) (Fig. 3C) along with two other NLP family TFs and a homolog of *TGA1*. While four TFs showed type-modulated response, only one was observed to be type-specific: *NLP7* (*Solyc08g082750*) which is down-regulated in epidermal cells.

**Figure 3.**
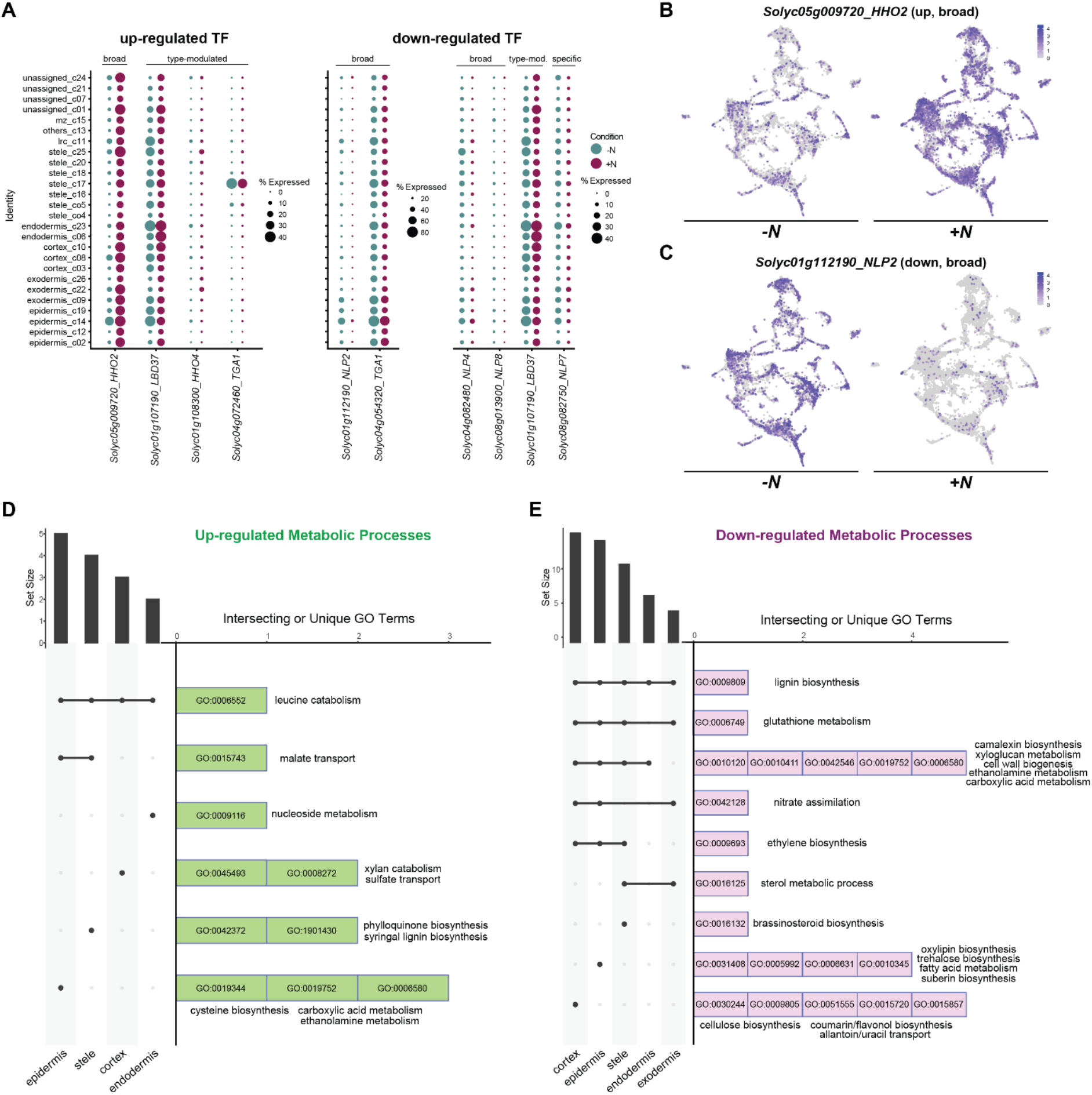
(A) Dot plots showing expression across the clusters for N-responsive DEGs within TF families associated with regulation of N response. (B-C) Maps of gene expression for exemplary N regulatory TFs with broad up- or down-regulation. Color represents scaled expression level. (D-E) Plots showing overlapping or unique GO terms enriched for metabolism-related processes in different cell/tissue level groupings.

The differentially expressed genes identified in different tissue and cell-type groupings (epidermis, stele, cortex, endodermis, or exodermis) were used to identify significantly enriched GO terms for those cell types, with a focus on GO terms related to metabolism (Fig. 3D-E). Based on enriched GO terms associated with up-regulated DEGs, the most broadly occurring change in metabolism is related to branched chain amino acid (leucine) catabolism, with genes encoding 3-methylcrotonyl-CoA carboxylase (*SlMCCA/Solyc09g065540* and *SlMCCB/Solyc01g108030*), isovaleryl-CoA-dehydrogenase (*SlIVD/Solyc11g069180*) and 3-Hydroxy-3-methylglutaryl-CoA lyase (*SlHMGL/Solyc01g080170*) showing increased expression across multiple cell types. Genes involved in malate transport were activated in the epidermis and stele; most notably a copy of a gene encoding aluminum-activated malate transporter 9 (*SlALMT9*/*Solyc05g009580*) was strongly induced in atrichoblasts. Among GO terms enriched in down-regulated DEGs, lignin biosynthesis genes showed a broad decrease in expression across all cell types tested, with the strongest enrichment in the cortex, where 12 lignin related-genes were down-regulated. Genes involved in xyloglucan metabolism and cell wall biogenesis were also down-regulated across most cell types (but not the exodermis); the strongest effect was again in the cortex, where 9 xyloglucan metabolic genes (mostly encoding xyloglucan endotransglucosylase/hydrolase family genes) were down-regulated.

Changes in hormone metabolism were also observed, with down-regulation of genes involved in producing ethylene through the ethylene-forming enzyme and 1-aminocyclopropane-1-carboxylate synthase 6 (*SlEFE/Solyc07g049550, SlACS6a/ Solyc12g056180*, and the stele-specific *SlACS6b/Solyc08g081535*) in the stele, cortex, and epidermis, and down-regulation of brassinosteroid biosynthesis (including *SlDWARF1/Solyc02g069490*) in the stele. The overall picture of dynamic processes in the root response to nitrate supply is a mixture of broadly distributed and more localized changes in metabolism and transport, particularly affecting energy resources, cell wall biosynthesis, and hormone signaling.

### A root single cell gene regulatory network reveals cell-type-specific circuits of nitrogen response

Gene regulatory networks (GRNs), which capture the web of TF-to-target interactions at a genome-wide scale, can be constructed from large transcriptomic datasets through regression-based approaches (Huynh-Thu et al. 2010). The adoption of single cell sequencing technology allows generation of large expression matrices suitable for generating GRNs (Badia-I-Mompel et al. 2023; Pratapa et al. 2020). Here, we utilized GRNBoost2 (Moerman et al. 2019) to generate a tomato root GRN across -N and +N conditions, which has the capability to identify TFs associated with cell type-specific regulatory networks of nitrogen response.

The resultant tomato root GRN has 977 TFs forming 454,483 predicted regulatory edges with 17,207 expressed genes. To identify TF regulators associated with gene regulation in a specific cell-type cluster, the predicted targets of each TF were compared with the top 100 marker genes of each cluster, and the significance of the overlap/enrichment was determined using hypergeometric distribution tests with multiple testing correction. Across the 26 root cell clusters, 730 TFs were significantly associated with regulation of cell type markers in at least one cluster. These TFs were hierarchically clustered based on the enrichment of cluster-specific marker genes among their predicted targets (Fig 4A). The clustering pattern of TFs generally recapitulates the cell-type assignments and relationships. For example, within the epidermis, the two clusters of atrichoblast cells (clusters 2 and 12) and the two trichoblast cells clusters (clusters 14 and 19) have strongly convergent patterns of enriched TFs within their specific epidermal sub-type, while a set of overlapping TFs also connect the trichoblast cells to the atrichoblasts, in line with a shared umbrella of epidermal cell identity. While similar connectivity is seen through the clusters of cortex, endodermal, and stele cells, the exodermal cells split: the cluster 22 and cluster 26 exodermal cells associated with active development of the exodermis in the root meristem have distinct TFs from the cluster 9 exodermal cells. Enrichment patterns for cluster 9 regulators show much more similarity to those for unassigned cluster 1 cells, which is in line with observations previously noted regarding their cluster type markers.

**Figure 4.**
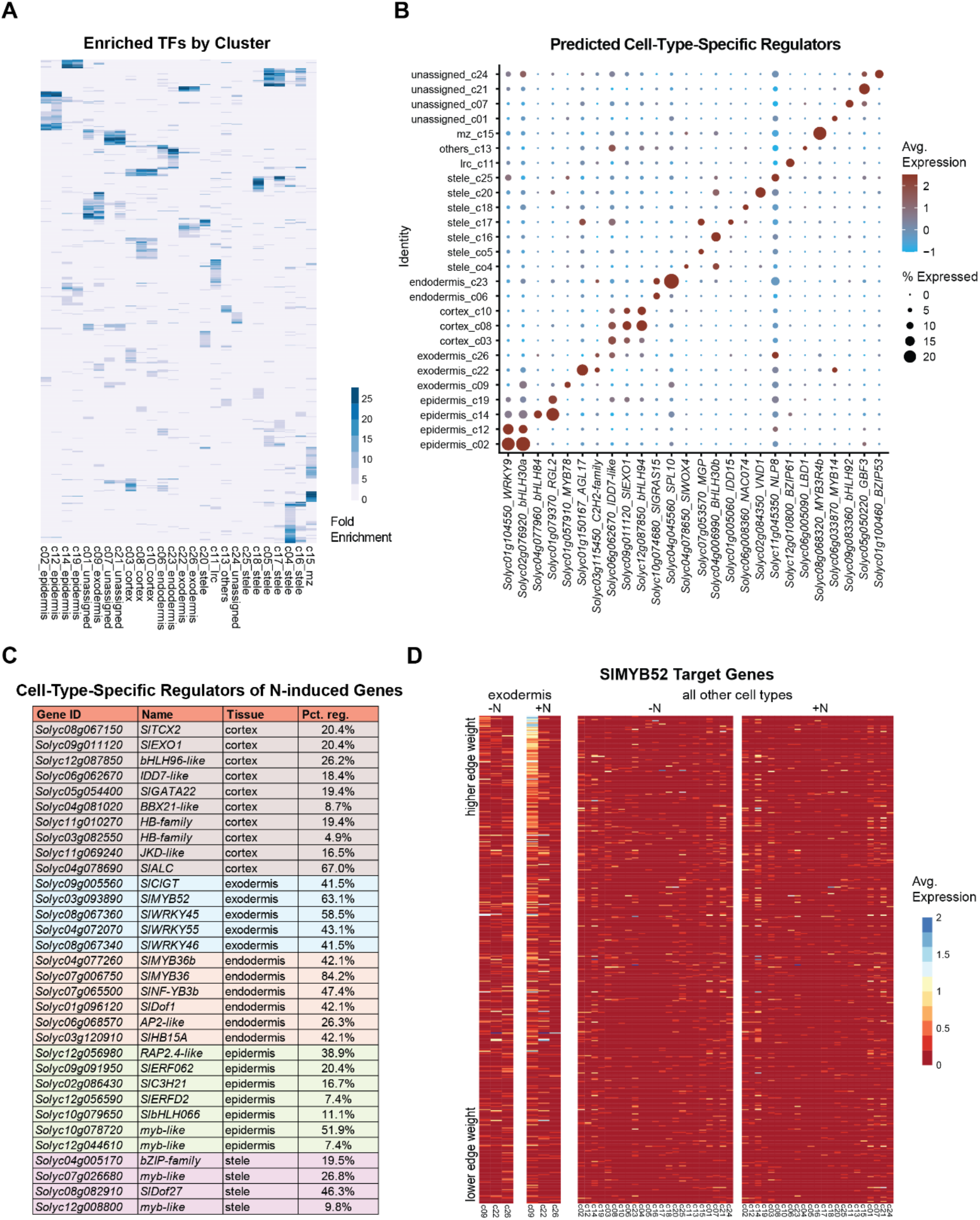
(A) Hierarchically clustered heatmap of fold enrichment of different TFs from the root GRN as measured by the relative proportion of their predicted targets that overlap with the top 100 marker genes from each cluster. (B) Dot plots showing expression of different predicted cell-type-specific regulators across combined data for each cluster. (C) Predicted regulators of cell-type-specific N-induced genes, with the percentage of the cell-type-specific up-regulated genes that are targets in the network for each TF. (D) Heat map of the expression of predicted targets of the exodermal cell-specific N response regulator SlMYB52.

From the sets of TFs predicted to regulate marker genes of each cluster (Fig. 4A), we further identified TFs with preferential expression in the cell-type cluster where their predicted target genes are also preferentially expressed, as these TFs are strong candidates for master regulators of biological processes associated with specific cell identities. Some prominent examples of such TFs meeting these criteria are plotted in Fig 4B. These predicted regulators include TFs previously known to be associated with a specific cell-type (S. Fig 10), as well as novel regulators. For example, in line with previously described plant cell type regulators, *SlWOX4* (*Solyc04g078650*) is the top predicted regulator of procambial markers (Ji et al. 2010), *SlPFA1* (*Solyc01g150101*) is the second ranked predicted regulator of markers in the cluster 5 pericycle cells (Ye Zhang et al. 2021), *SlVND1* (*Solyc02g084350*) and *SlVND7* (*Solyc11g018660*) are highly ranked predicted regulators of xylem markers (Kubo et al. 2005), and two *SlMYB3R4* genes (*Solyc11g071300, Solyc08g068320*) are highly associated with meristematic zone markers (Yang et al. 2021). Overall, these observations support that the GRN derived from single cell data is able to generate meaningful functional predictions.

To identify TFs important for the activation of transcriptional circuits involved in cell-type-specific N responses, the predicted targets of each TF were tested for significant enrichment against the type-specific N-induced gene sets (Fig 2A). Next, the identified TFs were further filtered by requiring that the TF is significantly upregulated by N in the same cell type in which it was predicted to regulate cell-type–specific N responses. This yielded a list of 32 strong candidate TFs for cell-type-specific regulation of N-induced gene activation (Fig. 4C). Interestingly, the list is largely devoid of any previously characterized regulators of N response which have been described at the organ or whole plant level. The regulated targets of these TFs are also generally not enriched in sentinel genes canonically involved in nitrogen metabolism (Para et al. 2014; Julian, Patrick, and Li 2023), except for an AP2-like TF (*Solyc06g068570*) acting in the endodermis, which is predicted to regulate the expression of 3 transporters and an asparaginase gene.

Due to its relatively high proportion of type-specific N-induced genes (Fig 2A), we focused on the exodermal cells to study the cell-type-specific TF regulators of N response, which were all specifically expressed in cluster 9. Among these, particularly high enrichment was noted for *SlCIGT* (*Solyc09g005560*), a homolog of the *Solanum habrochaites* stress-inducible Trihelix transcription factor *ShCIGT* (C. Yu et al. 2018), while MYB family transcription factor *SlMYB52* (*Solyc03g093890*) was predicted to regulate the largest portion of type-specific genes, forming regulatory edges to 41 of the 65 genes type-specifically up-regulated by N in cluster 9 cells (63.1%; Fig 4C).

The network of 451 predicted target genes for SlMYB52 was examined for expression levels across the root cell clusters in -N and +N condition (Fig 4D). As expected, these target genes show preferential expression specifically in cluster 9 cells. Many targets showed strong induction in cluster 9 in response to N, with stronger transcriptional induction associated with stronger predicted edge weight in the regulatory network. Examining the exodermal-specific N-induced genes regulated by SlMYB52, we noted multiple genes (S. Fig 11) encoding proteins associated with benzenoid metabolism (Huang et al. 2022; Boersma et al. 2022; Huang, Lee, and Dudareva 2025), including cinnamoyl-CoA ligase (*SlCNL1, Solyc02g081360*), cinnamoyl-CoA hydratase-dehydrogenase (*SlCHD1, Solyc07g019670*), 3-ketoacyl-CoA thiolase, (*SlKAT1, Solyc09g061840*), a benzaldehyde synthase α-like gene (*SlBSα-like, Solyc10g080900*), and two loci encoding genes for benzoyl-CoA:benzyl alcohol benzoyl transferase (*SlBEBTa, Solyc07g049645*; *SlBEBTb, Solyc07g049655*).

## DISCUSSION

Our single cell study of tomato roots exposed to different N environment painted a comprehensive picture of how individual cell-types coordinate the organ level response to nitrogen availability (Fig. 1). A significant overlap was observed between the differentially expressed genes (DEGs) identified from bulk root tissues in our previous study (Julian, Patrick, and Li 2023), and those detected in the current single-cell analysis (Fig. S9). Because both experiments were performed under identical N treatments and at the same time point, this overlap validates the robustness of the single-cell dataset. Notably, the overlapping genes are dominantly regulated by N across multiple tissue types. Meanwhile, the single-cell analysis revealed a large set of N-responsive genes that exhibit strong cell-type specificity (Fig. 2), which is in line with the expectation that scRNA-Seq allows more insight into tissue- and cell-type-specific transcriptional dynamics that were masked in bulk, organ-level transcriptomic analyses, thus providing higher spatial resolution. It is also worth noting that that bulk tissue RNA-seq measures steady state mRNA levels, while scRNA-Seq experiment measures RNA levels in nuclei, which is enriched with recently transcribed nascent RNA, therefore providing additional sensitivity to transcriptional dynamics.

Examining the enrichment of biological processes among DEGs across cell types (Fig. 3) provided additional insights into how different cell layers respond to N changes. In response to N supply, there is a prominent down-regulation of genes involved in cell wall and lignin biosynthesis across different cell layers. This is in agreement with a meta-analysis showing that often N availability and lignin biosynthesis are negatively correlated (Peng et al. 2024). This could reflect that during N deficiency, cell wall and lignin are strengthened to facilitate and maximize water/nutrient uptake and transport to allow plants to survive the adverse conditions. The observed up-regulation of leucine catabolic pathways across multiple root cell types in response to nitrogen resupply is consistent with our previous report showing a reduction in leucine content following nitrogen supply (Y. Li et al. 2020). One possible explanation is that leucine catabolism primarily serves as a source of metabolic precursors, providing carbon skeletons to the TCA cycle to support enhanced primary metabolism and growth following N resupply. Leucine is the most abundant amino acid in Arabidopsis, and its degradation yields one of the highest energy outputs among amino acids (Hildebrandt et al. 2015). One the other hand, in mammalian systems, leucine serves as a key molecular indicator of nutrient status by activating the mTOR pathway to promote protein synthesis (Dodd and Tee 2012). The TOR signaling pathway is conserved in plants (Shi, Wu, and Sheen 2018), and emerging evidence suggests that leucine or other branched-chain amino acids (BCAAs) may also contribute to TOR activation in plants (P. Cao et al. 2019). In plants, nitrogen availability is known to activate TOR signaling (Yanlin Liu et al. 2021). However, this presents an apparent paradox: TOR activity is enhanced under N-sufficient conditions, and BCAAs such as leucine are proposed activators of TOR, yet leucine is actively degraded upon nitrogen resupply. Resolving whether leucine plays a direct signaling role in plant TOR activation, or instead functions predominantly as a metabolic substrate, will require further investigation.

Focusing on the cell-type-specific and dynamic N responses, we identified multiple novel TF regulators that have been under-explored in previous bulk-tissue level studies (Fig. 4). Interestingly, the predicted targets of these TFs are generally not directly involved in N metabolism. In one prominent case with cell-specific regulation by SlMYB52, the predicted targets are enriched in specialized metabolism pathways. This is consistent with the concept that primary metabolism is a core function performed across many cell-types while specialized metabolism can be more specific to tissue types (Y. Li, Grotewold, and Dudareva 2024). These findings point to previously unrecognized layers of N-responsive regulation that extend beyond the core N metabolic pathways. For example, SlMYB52 is predicted to activate multiple genes involved in benzenoid metabolism in response to N. Collectively, the induced enzymes SlCNL1, SlCHD1, and SlKAT1 can use trans-cinnamic acid (produced from phenylalanine by PAL - phenylalanine ammonia lyase) to generate benzoyl-CoA. This can be combined with benzyl alcohol (possibly produced downstream of benzaldehyde synthase) by the SlBEBT enzymes to produce benzylbenzoate. This indicated that increased nitrate levels induce production of benzylbenzoate in this cell population, though the reason was not clear. However, recently it has been elucidated that benzylbenzoate produced through this set of enzymes is connected to the biosynthesis of salicylic acid (SA) through the action of two additional enzymes which act on the benzylbenzoate intermediate (Y. Wang et al. 2025; Yanan Liu et al. 2025; K. Li et al. 2025; Zhu et al. 2025; Ma et al. 2025). Checking our data, we found that genes for these enzymes, benzylbenzoate hydroxylase (*SlBBH, Solyc03g122350*) and benzylsalicylate esterase (*SlBSE, Solyc02g069800*) are also predicted to be regulated by SlMYB52 and induced in response to N in cluster 9 cells (S. Fig 11). Therefore, among the set of 41 genes predicted to be regulated by SlMYB52 in a cell-type-specific manner during N response, the entire phenylalanine-dependent SA biosynthesis network was captured, highlighting the power of the GRN-based approach using single cell data. The specific role of SA in nitrogen response will require further investigation.

