## SupplementalFigures for "Every Cell Counts: Tomato Root Responses to Nitrogen at Single-Cell Resolution"

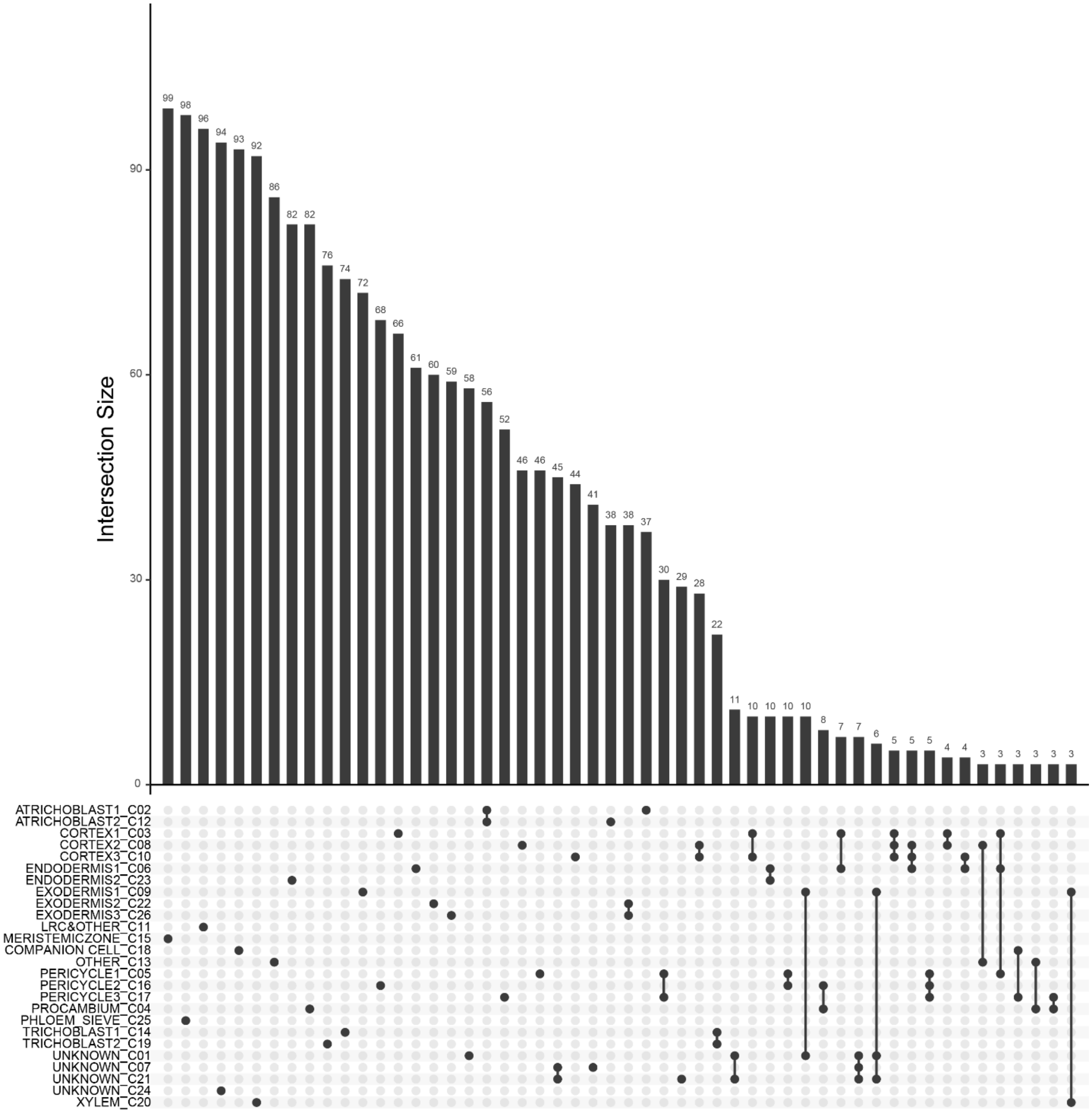


Supplemental Figure 1. Intersections of the top 100 marker genes from each cluster aid identification of closely related cell types.

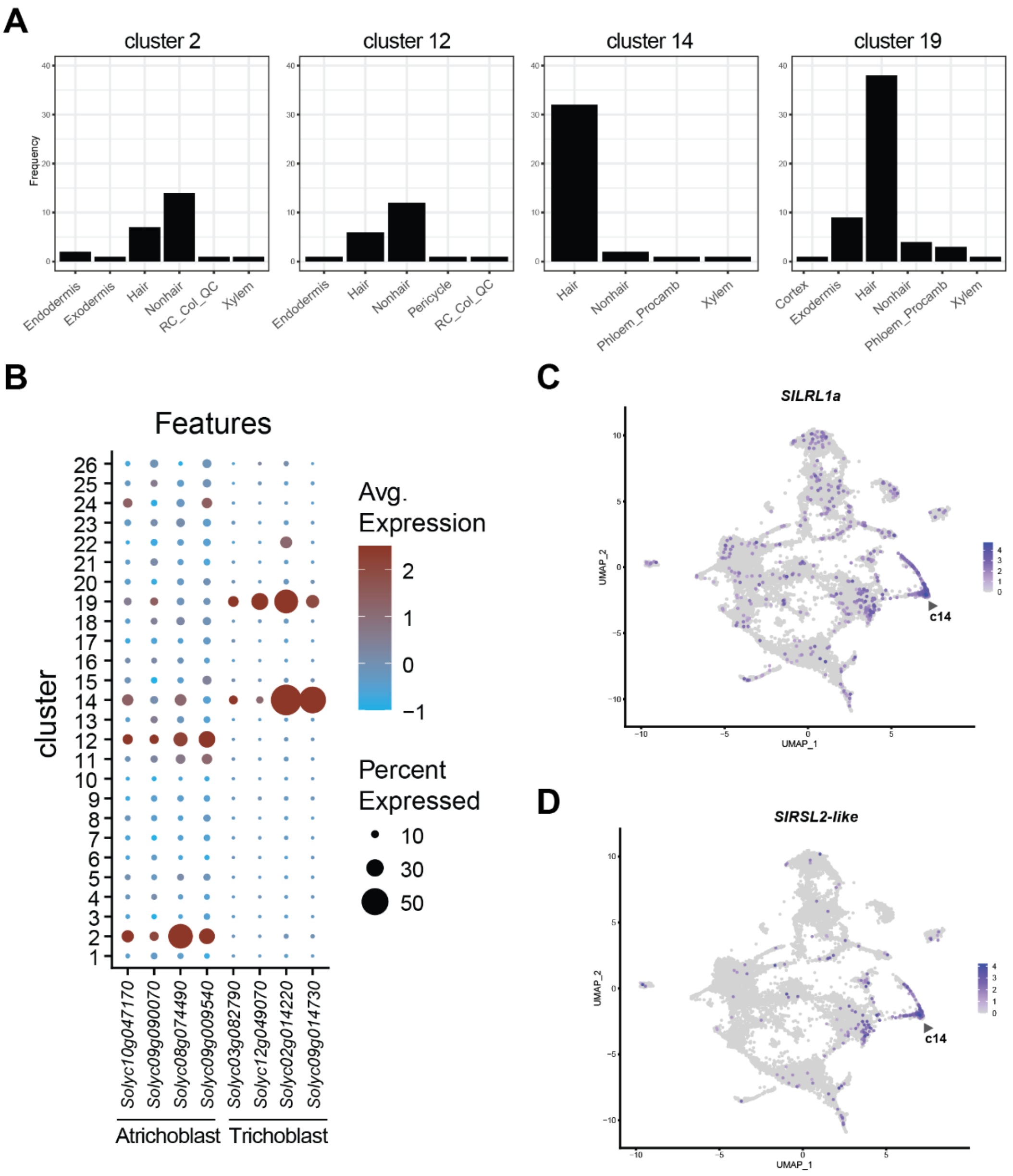


Supplemental Figure 2

(A) Categorization of the top 100 marker genes in each cluster according to cross-species orthologous marker groups for roots (Chau et al. 2025) shows plurality assignment of clusters 2, 12 to nonhair cell-types and clusters 14, 19 to hair cell-types. (B) Root epidermal marker genes previously identified from tomato (Cantó-Pastor et al. 2024) or cross-species orthologous marker groups (Chau et al. 2025). Atrichoblast markers *SlSULTR1;3* (*Solyc10g047170*), *SlPHT1;4* (*Solyc09g090070*), *Nucleotide-diphospho-sugar transferase family protein* (*Solyc08g074490*), *alpha/beta-Hydrolases superfamily protein* (*Solyc09g009540*). Trichoblast markers *SlDif54* (*Solyc03g082790*), *SlEXT1* (*Solyc12g049070*), *SlGH9C1* (*Solyc02g014220*), *SlWAK3* (*Solyc09g014730*). (C-D) Certain root hair-specific regulatory genes, e.g. *SlLRL1a (Solyc12g010170)* and *SlRSL2-like (Solyc12g088380)*, are strongly enriched in cluster 14 expression.

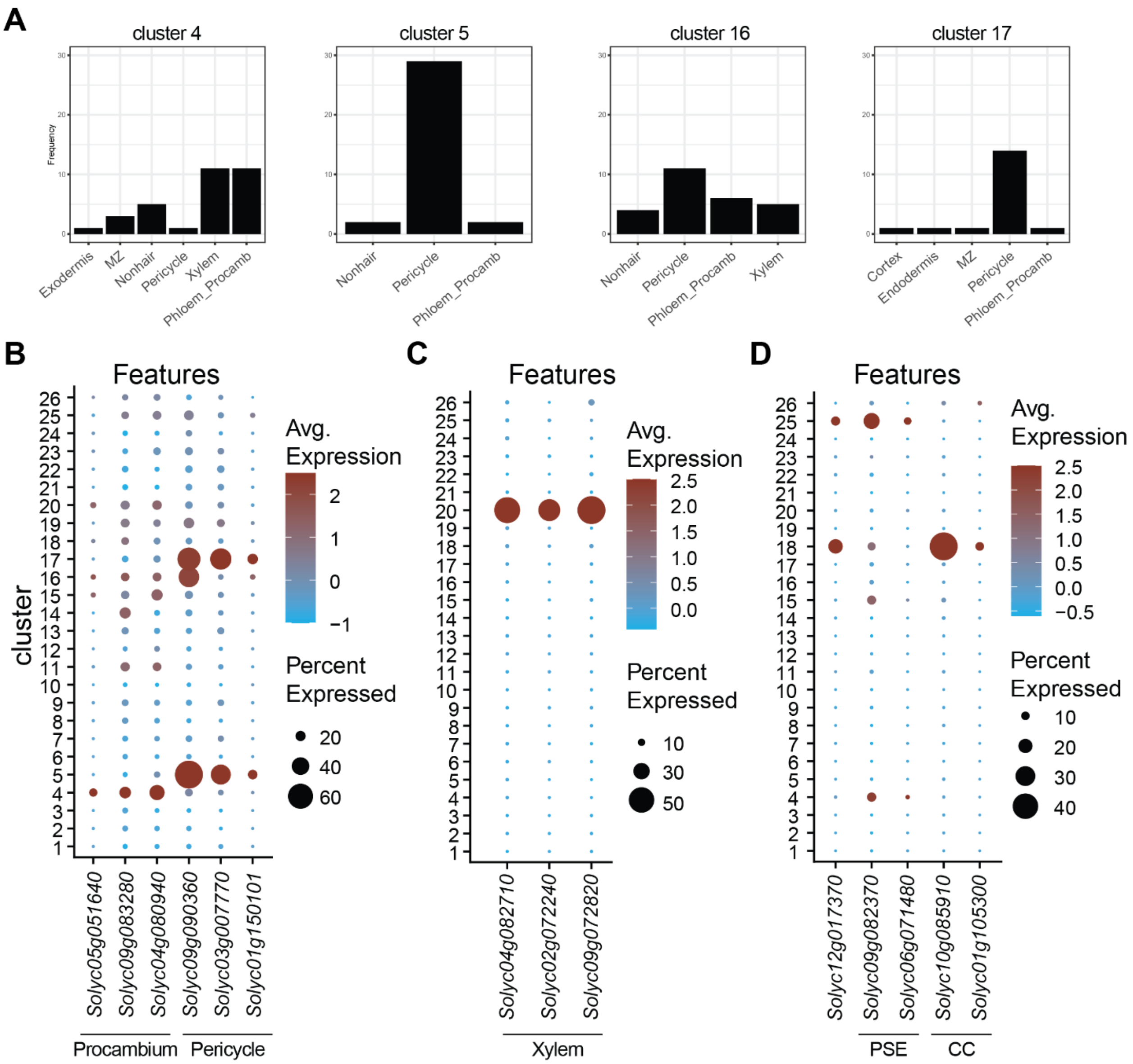


Supplemental Figure 3

(A) Categorization of the top 100 marker genes in each cluster according to cross-species orthologous marker groups for roots (Chau et al. 2025) finds cluster 4 most similar to phloem/procrambium and xylem assignments, while clusters 5, 16, 17 have majority or plurality pericycle assignment. (B) Root procambium and pericycle marker genes previously identified from tomato (Cantó-Pastor et al. 2024) or cross-species orthologous marker groups (Chau et al. 2025). Procambium markers *SlPXY* (*Solyc05g051640*), *SlSHY2* (*Solyc09g083280*), *SlWAT1* (*Solyc04g080940*) and pericycle markers *SlPHO1* (*Solyc09g090360*), *SlSLAH3* (*Solyc03g007770*), *SlPFA1* (*Solyc01g150101*). (C) Root xylem marker genes previously identified from tomato (Canto-Pastor et al): *SlXCP2* (*Solyc04g082710*), *SlIRX1* (*Solyc02g072240*), *SlIRX5* (*Solyc09g072820*). (D) Phloem cell markers include the developmental regulator *SlAPL* (*Solyc12g017370*) and marker genes (Otero et al. 2022; Miyashima et al. 2019) associated with phloem sieve elements (PSE) *SlCVP2* (*Solyc09g082370*), *SlHCA2* (*Solyc06g071480*), or companion cells (CC) *SlNAKR1* (*Solyc10g085910*), *SlMC3* (*Solyc01g105300*).

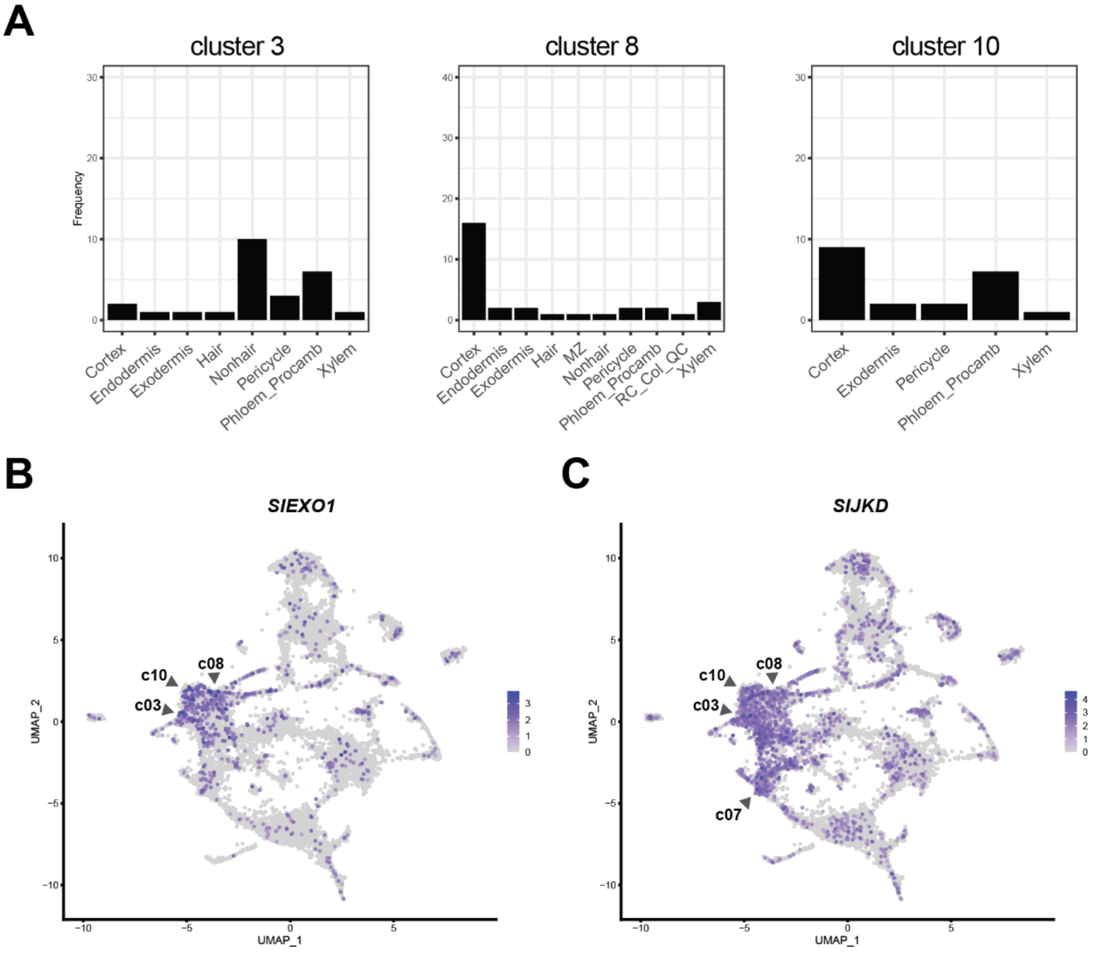


Supplemental Figure 4. (A) Categorization of the top 100 marker genes in each cluster according to cross-species orthologous marker groups for roots (Chau et al. 2025) show plurality assignment to cortex for clusters 8, 10 while cluster 3 markers are associated with nonhair epidermal cells and phloem/procambium across orthologous marker groups. (B-C) The inner cortical cell marker (Manzano et al. 2025) *SlEXO1* (*Solyc09g011120*) is expressed in clusters 3, 8, and 10, while the cortical cell associated transcription factor (Hassan, Scheres, and Blilou 2010; Liang et al. 2024) *SlJKD* (*Solyc10g084180*) is expressed strongly in these clusters as well as 7.

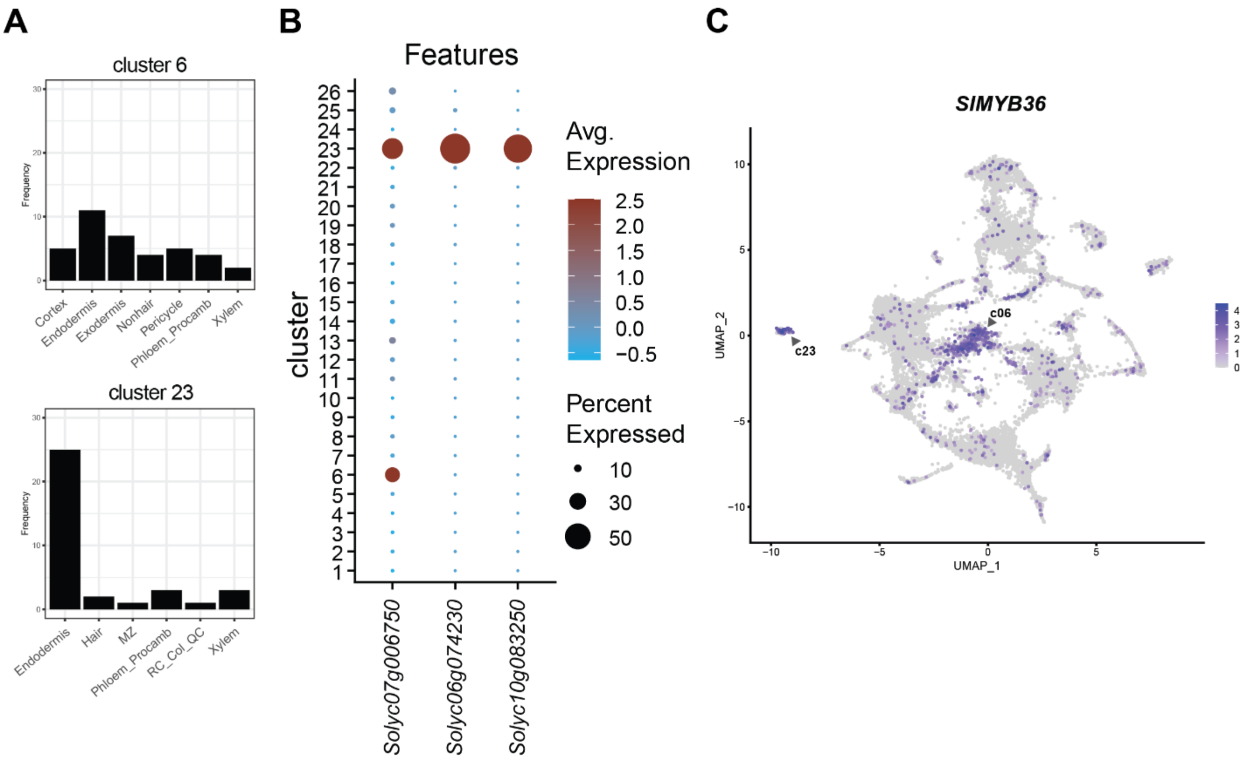


Supplemental Figure 5. (A) Categorization of the top 100 marker genes in each cluster according to cross-species orthologous marker groups for roots (Chau et al. 2025) finds a strong majority assignment for endodermis in cluster 23 and a plurality assignment for cluster 6. (B-C) Endodermal marker genes from previous work in tomato (Cantó-Pastor et al. 2024). Casparian strip associated genes *SlCASP1* (S*olyc06g074230*), *SlCASP2* (*Solyc10g083250*), are expressed predominantly in cluster 23, while the master regulator of endodermal cell identity *SlMYB36* (*Solyc07g006750*) is expressed across both clusters 6 and 23.

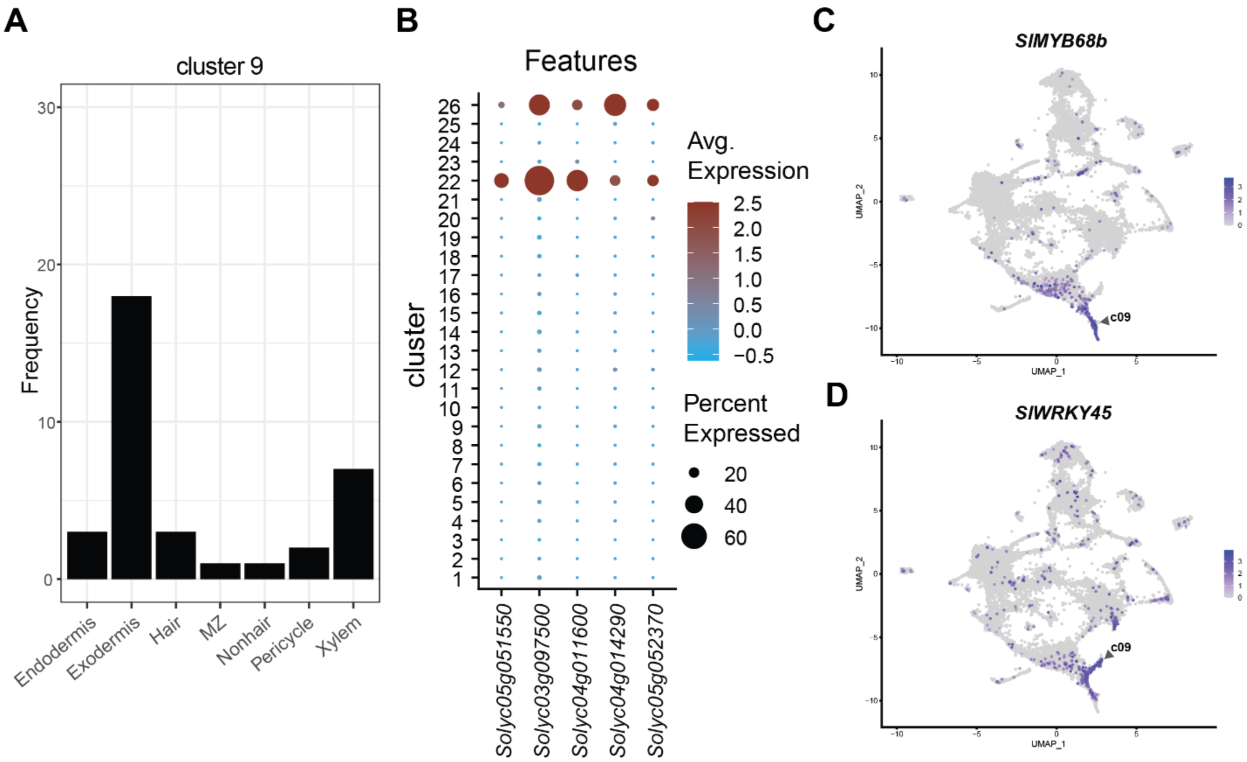


Supplemental Figure 6. (A) Categorization of the top 100 marker genes in each cluster according to cross-species orthologous marker groups for roots (Chau et al. 2025) shows a majority assignment to exodermis for cluster 9. Clusters 22, 24 markers did not have significant overlap with orthologous marker groups. (B) Previously identified exodermal marker genes from the root tip (Cantó-Pastor et al. 2024): the TF *SlMYB92* (*Solyc05g051550*) which is associated with developing exodermis is expressed in cluster 22 while other markers *SlASFT* (*Solyc03g097500*), *SlGAPT5* (*Solyc04g011600*) are expressed in clusters 22 and 26. Genes associated with the cell wall such as *SlPELPK* (*Solyc04g014290*), *SlLAC12-like* (*Solyc05g052370*) are enriched in cluster 26 while also being expressed in cluster 22. (C-D) The two branches of cluster 9 have partly diverging sets of markers, for example the distributions of TFs *SlMYB68b* (*Solyc06g074910*) and *SlWRKY45* (*Solyc08g067360*).

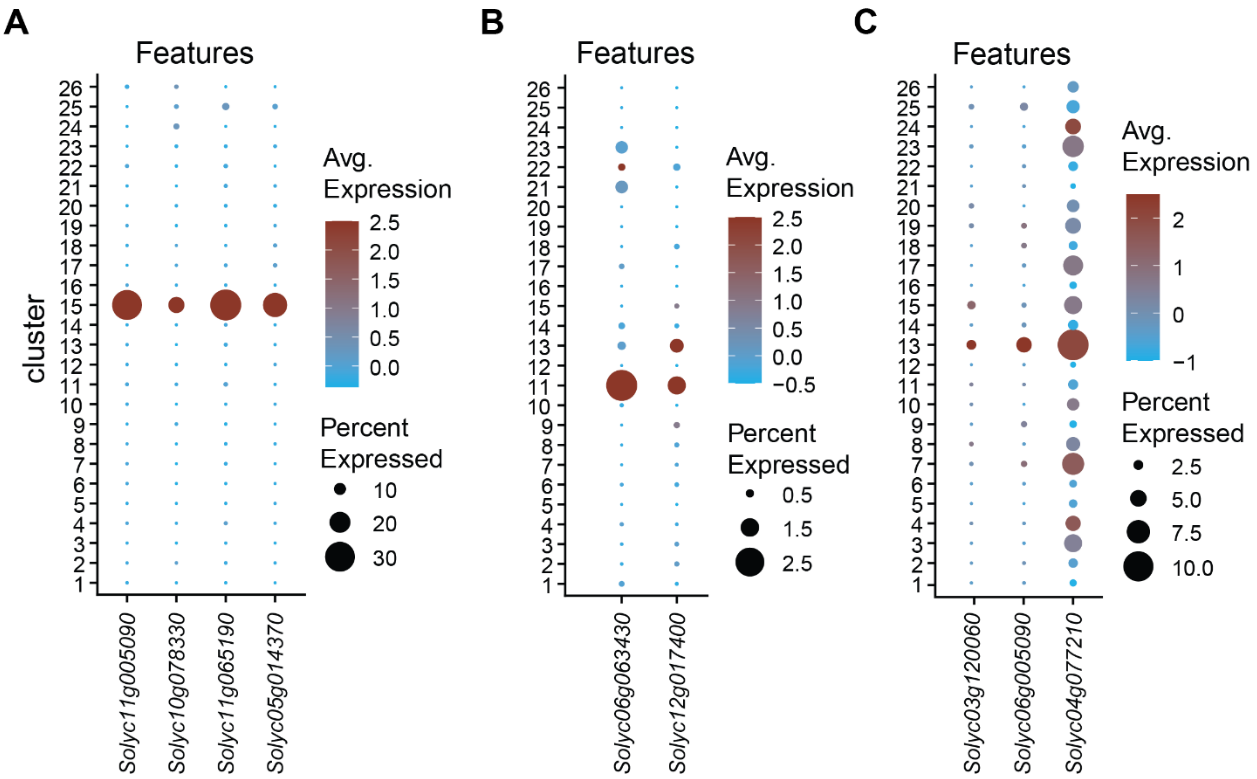


Supplemental Figure 7. (A) Meristemic zone markers from tomato (Cantó-Pastor et al. 2024) or orthologous gene markers (Chau et al. 2025) with supporting observations in Arabidopsis (Komaki and Schnittger 2017; Criqui et al. 2002): *SlCYCA1;1* (*Solyc11g005090*), *SlCYCB1;1* (*Solyc10g078330*), *SlUBC19* (*Solyc11g065190*), *SlMAD2* (*Solyc05g014370*). (B) The markers associated with emerging lateral roots (Bennett et al. 2010): *SlBRN2* (*Solyc06g063430*), *SlSMB* (*Solyc12g017400*). (C) Markers associated with cluster 13 cells: *SlBAS1* (*Solyc03g120060*) expressed in the root cap (Chaiwanon and Wang 2015), *SlLBD1* (*Solyc06g005090*) and *SlKNAT1* (*Solyc04g077210*) associated with secondary growth (Woerlen et al. 2017; Ye et al. 2021).

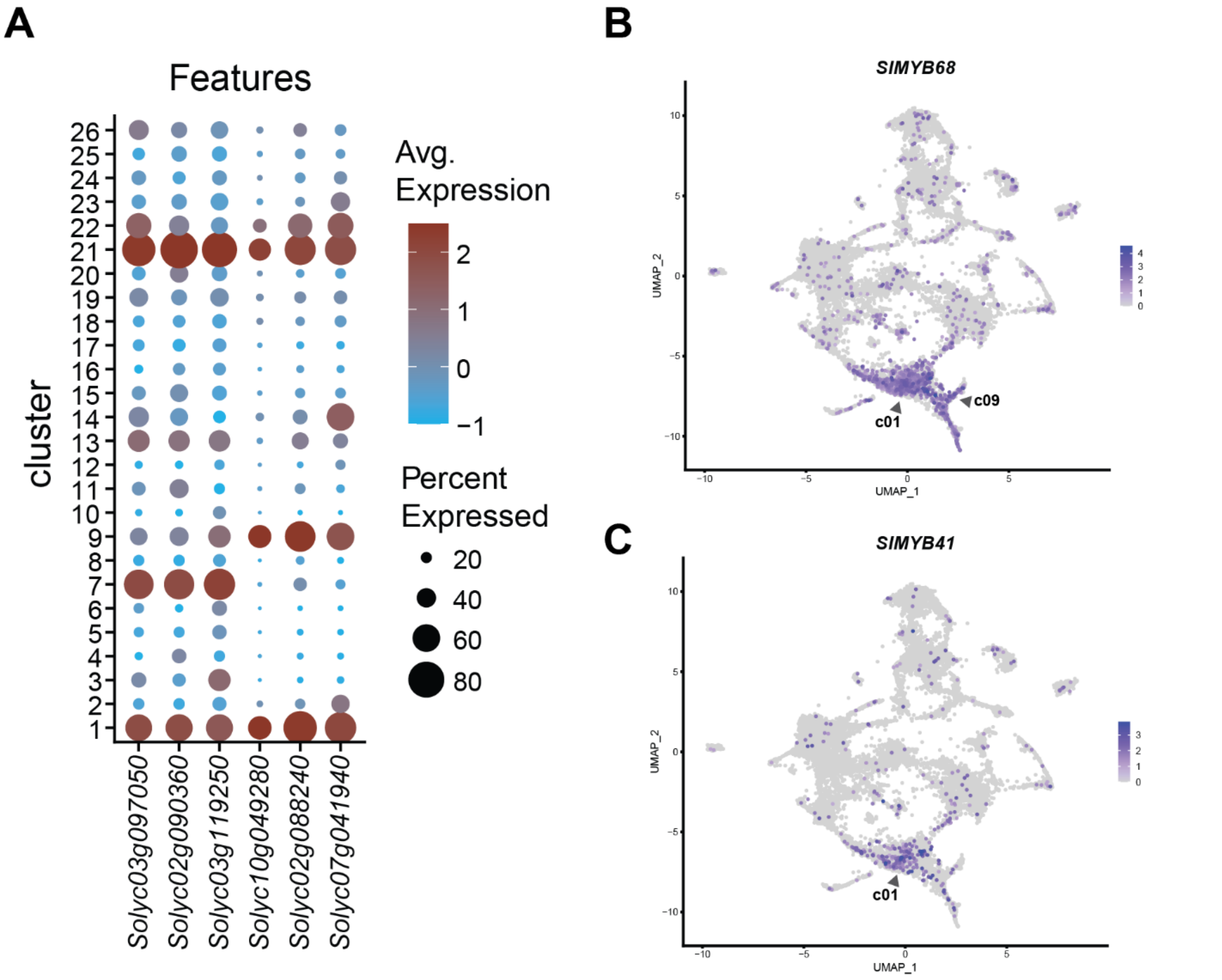


Supplemental Figure 8. (A) Shared gene markers for clusters 1-7-21 *SlCSLD* (*Solyc03g097050*) *SlSKS17* (*Solyc02g090360*) *SlCBP60* (*Solyc03g119250*); shared gene markers for clusters 1-9-21 *SlNAT* (*Solyc10g049280*) *SlPHO1-H* (*Solyc02g088240*) *SlRIPK* (*Solyc07g041940*). (B-C) *SlMYB68* (*Solyc11g069030*), a member of the group of TFs associated with suberization (Manzano et al. 2025; Kraska et al. 2025) is expressed in clusters 1 and 9, while the recently identified exodermal regulator of suberization *SlMYB41* (*Solyc02g079280*) is expressed in cluster 1.

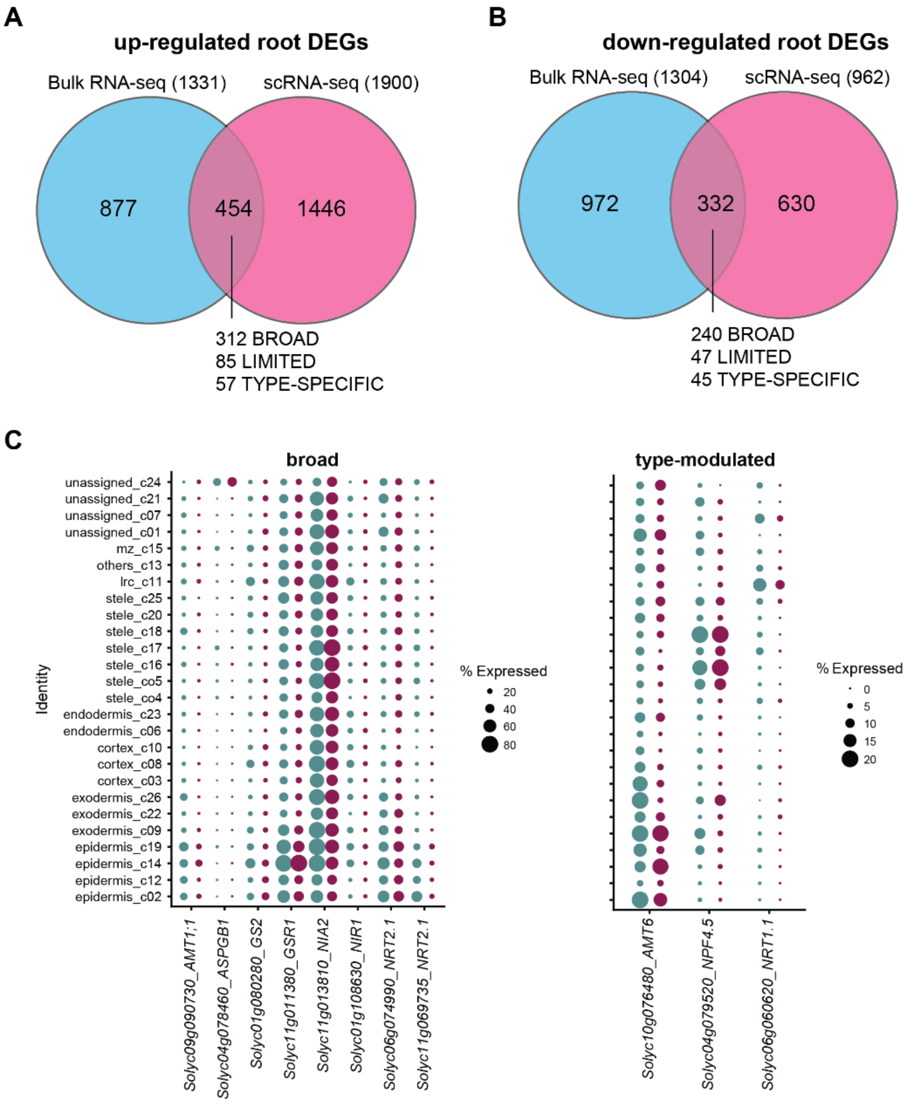


Supplemental Figure 9. (A-B) Overlaps between previous bulk root tissue RNA-Seq in a previous +N treatment experiment with similar conditions (Julian, Patrick, and Li 2023) and DEGs from single cell RNA-Seq. The overlapping DEGs are broken out by category. (C) Dot plots showing expression across the clusters for down-regulated N-responsive DEGs involved in N metabolism and transport across broad and type-modulated categories.

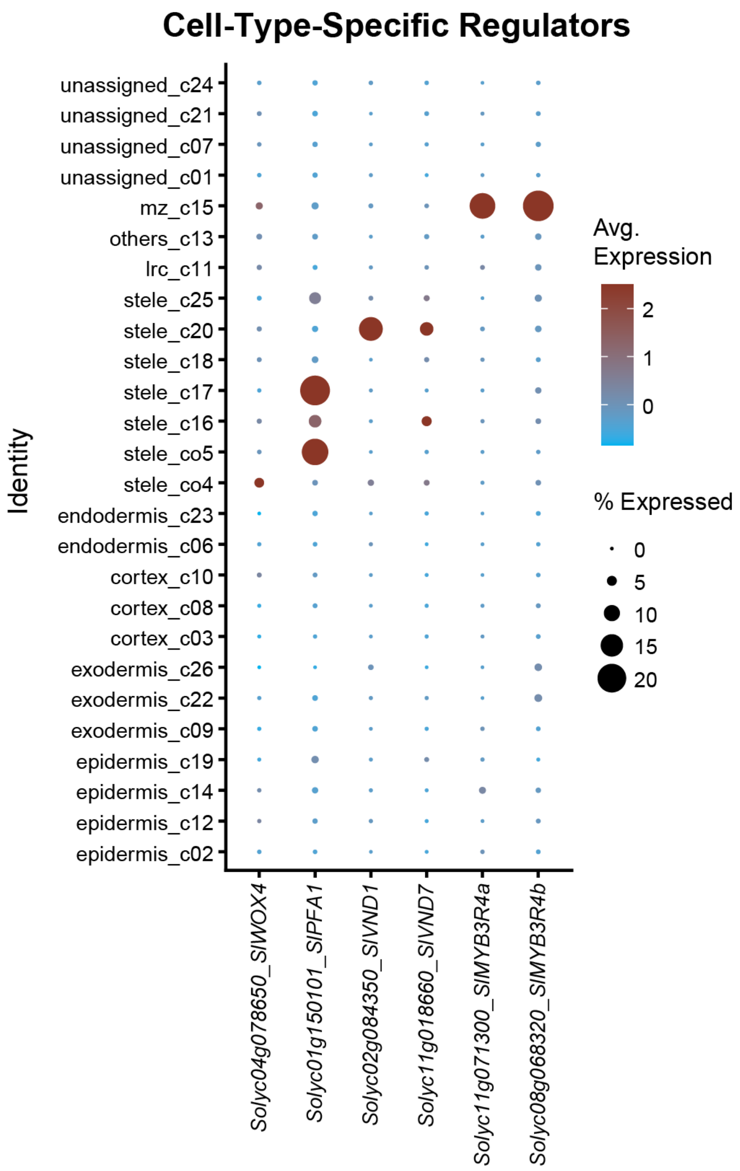


Supplementary Figure 10. Some examples of predicted transcriptional regulators from the tomato root GRN that comport with previously described cell-type-specific roles based on the expression patterns of the TFs and their predicted regulatory targets.

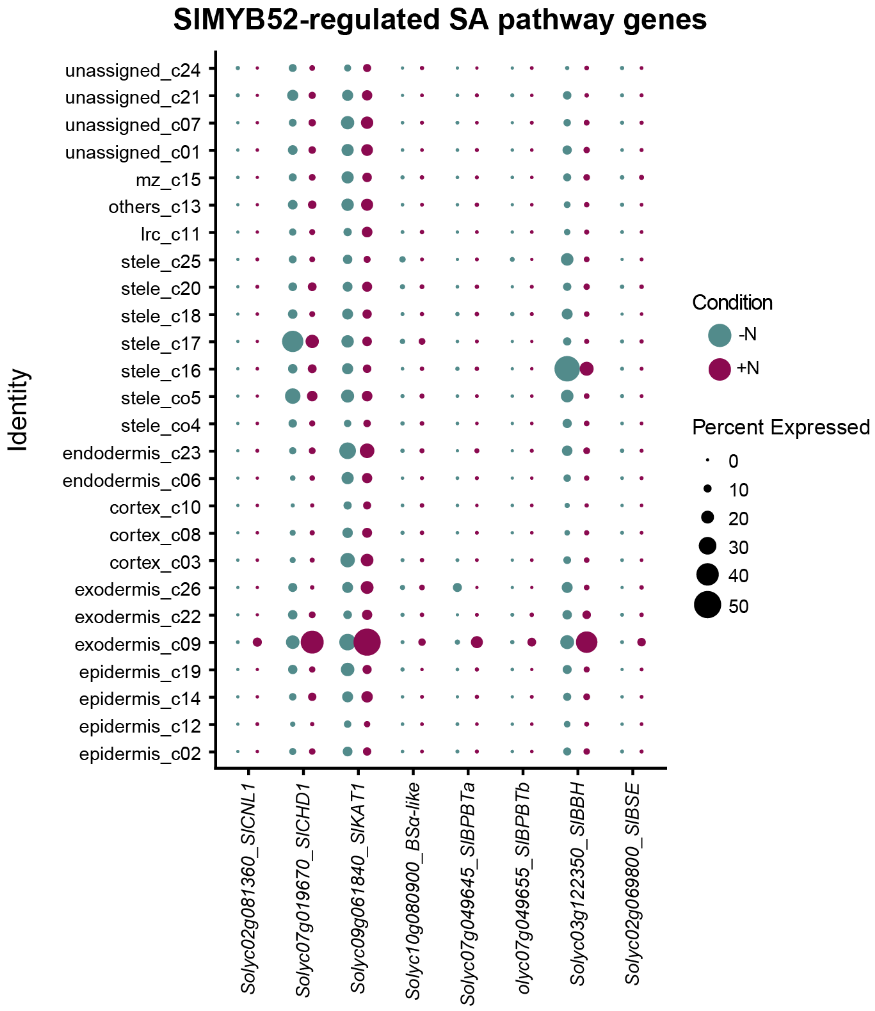


Supplementary Figure 11. Expression patterns of genes involved in benzenoid metabolism and the phenylalanine-dependent salicylic acid (SA) pathway in the root single cell data. These genes are all predicted to be collectively co-regulated by SlMYB52 and type-specifically induced in cluster 9 exodermal cells in response to N supply.

Cantó-Pastor, Alex, Kaisa Kajala, Lidor Shaar-Moshe, Concepción Manzano, Prakash Timilsena, Damien De Bellis, Sharon Gray, et al. 2024. "A Suberized Exodermis Is Required for Tomato Drought Tolerance." *Nature Plants* 10 (1): 118–30.

Chau, Tran N., Prakash Raj Timilsena, Sai Pavan Bathala, Sanchari Kundu, Bastiaan O. R. Bargmann, and Song Li. 2025. "Orthologous Marker Groups Reveal Broad Cell Identity Conservation across Plant Single-Cell Transcriptomes." *Nature Communications* 16 (1): 201.

Criqui, Marie Claire, Janice de Almeida Engler, Alain Camasses, Arnaud Capron, Yves Parmentier, Dirk Inzé, and Pascal Genschik. 2002. "Molecular Characterization of Plant Ubiquitin-Conjugating Enzymes Belonging to the UbcP4/E2-C/UBCx/UbcH10 Gene Family." *Plant Physiology* 130 (3): 1230–40.

Komaki, Shinichiro, and Arp Schnittger. 2017. "The Spindle Assembly Checkpoint in Arabidopsis Is Rapidly Shut off during Severe Stress." *Developmental Cell* 43 (2): 172-185.e5.

Miyashima, Shunsuke, Pawel Roszak, Iris Sevilem, Koichi Toyokura, Bernhard Blob, Jung-Ok Heo, Nathan Mellor, et al. 2019. "Mobile PEAR Transcription Factors Integrate Positional Cues to Prime Cambial Growth." *Nature* 565 (7740): 490–94.

Otero, Sofia, Iris Gildea, Pawel Roszak, Yipeng Lu, Valerio Di Vittori, Matthieu Bourdon, Lothar Kalmbach, et al. 2022. "A Root Phloem Pole Cell Atlas Reveals Common Transcriptional States in Protophloem-Adjacent Cells." *Nature Plants* 8 (8): 954–70.
